# Seasonal Variations and Challenges in Estimating Populations and Identifying Species of Korean Ungulates Using Drone-Derived Thermal Orthomosaic Maps

**DOI:** 10.1101/2025.01.23.633323

**Authors:** Jinhwi Kim, Donggul Woo

## Abstract

Thermal drones offer significant time and labor savings in estimating wild ungulate populations. Among available methods, Thermal Infrared (TIR) orthomosaic maps have shown potential for animal counting and population estimation. In this study, we assessed the feasibility of counting Korean ungulates in March and June using TIR and RGB orthomosaic maps, aiming to compare population estimates across season and identify the appropriate Ground Sampling Distance (GSD). The average estimated population was 25.3 ± 4.2 individuals (range: 18–29) in March and 28.2 ± 9.1 individuals (range: 14–40) in June. The average counting error was 4.1 (95% CI: 1.3–6.9) across the orthomosaic maps, with no substantial difference between March and June (P = 0.75). Counting errors increased when RGB GSD exceeded 2.5 cm/pix and TIR GSD exceeded 7.5 cm/pix, suggesting that these thresholds may guide future GSD selection for more accurate population estimates. We measured 149 thermal marks for species identification across three ungulate species. MANOVA revealed strong effects of species (P < 0.001) and posture (P < 0.001) on body measurements, but variations in posture limited the effectiveness of thermal mark measurements for species identification. This issue was especially prominent in lying positions, where species overlap occurred. Further data collection on thermal measurements from TIR orthomosaic maps, particularly from a perpendicular angle and across diverse environments and postures, could improve the accuracy of thermal methods for species identification.

## 1. Introduction

Accurate population estimates of endangered wildlife are essential for ecological research and the development of effective conservation plans. However, this process is complex and requires substantial costs, time, and labor(Dundas et al., 2021). Traditional methods for wildlife population estimation include direct field observations by surveyors along predetermined routes, inference based on signs of presence, and long-term monitoring using remote camera traps. These methods, however, often face challenges related to underestimation, particularly when populations are small or detection is difficult(Cukor et al., 2022; Gu and Swihart, 2004; Tóth and Katona, 2024). Studies have shown that traditional survey techniques can lead to systematic biases in population estimates, underscoring the need for improved methodologies that incorporate advanced monitoring technologies (Amos et al., 2014; Samuel et al., 1992; Willson et al., 2011; Zhang et al., 2021).

To address these challenges and enhance monitoring capabilities, unmanned aerial vehicles (drones) have emerged as a promising alternative. Recent advancements in battery performance have extended drone flight times, positioning them as viable tools to complement or even replace traditional field surveys(Dundas et al., 2021; Povlsen et al., 2023). Previous drone-based wildlife studies typically involve flying along predetermined routes to capture images and videos, which are then used to count individuals from an aerial perspective. This approach has been successfully applied to various mammal species, including white-tailed deer(Beaver et al., 2020; Davis, 2021.), deer species(Witczuk et al., 2018), moose(Mayer et al., 2024), and hippopotamuses(Inman et al., 2019). Compared to ground-based surveys, drone-based methods have demonstrated increased efficiency by identifying more individuals, reducing survey time, and covering larger areas(Beaver et al., 2020; Davis, 2021; Dundas et al., 2021; Mayer et al., 2024).

Along with the advancements in camera technology, it is now possible to create high-resolution orthomosaic maps. These maps are generated by accurately correcting and stitching together multiple aerial or satellite images, providing a true-scale representation of the terrain without distortion. Orthomosaic maps facilitate detailed assessments of wildlife habitats and movement paths, and have proven valuable for estimating population numbers in specific areas. Various studies utilizing orthomosaic maps include those on Galápagos iguanas(Varela-Jaramillo et al., 2023), bat colonies(McCarthy et al., 2022), Japanese macaque(Mirka et al., 2022), southern giant petrels(Fudala and Bialik, 2022), muskrat(Greenhorn et al., 2024), and Nile crocodile(Ezat et al., 2018).

Despite these advancements, forest-dwelling ungulates often remain concealed under dense vegetation, making population estimates labor-intensive and challenging. Studies by Amos et al.(Amos et al., 2014) and Pena et al.,(Peña-Carmona et al., 2024) highlight systematic underestimations in direct counts and density estimates of mountain ungulate populations. This issue is particularly relevant when considering the population estimates of Korean ungulate species, which have predominantly relied on non-invasive DNA analysis from fecal samples(Jang et al., 2020), and transect-line censuses of fecal deposits(Kim et al., 2011). The limitations of these approaches necessitate the exploration of more effective monitoring techniques tailored to the unique challenges presented by these species in their natural habitats.

This study aimed to evaluate the effectiveness of using a thermal drone to estimate the population of three Korean ungulate species in captivity, employing Thermal Infrared (TIR) orthomosaic maps. We compared the animal counts from these maps, generated under varying flight conditions, to identify the optimal Ground Sample Distance (GSD) for accurate population estimation and efficient drone operation. Additionally, we measured the thermal signature body sizes of the ungulate species to explore the feasibility of species identification using TIR orthomosaic maps.

## 2. Materials and Methods

### 2.1. Study site

We conducted 17 flights at the deer park within National Institute of Ecology, Seocheon, South Korea. The park houses three species of Korean ungulate: Water deer(*Hydropotes inermis*), Roe deer(*Capreolus pygargu*), and Long-tailed goral(*Naemorhedus caudatus*), in a 20,227m² fenced area for exhibition purposes. This area is predominantly flat and includes various structures such as trees, rock mounts, streams, swamps, three roofed feeding stations, and six net feeding stations (**Figure 1**).

**Figure 1.**
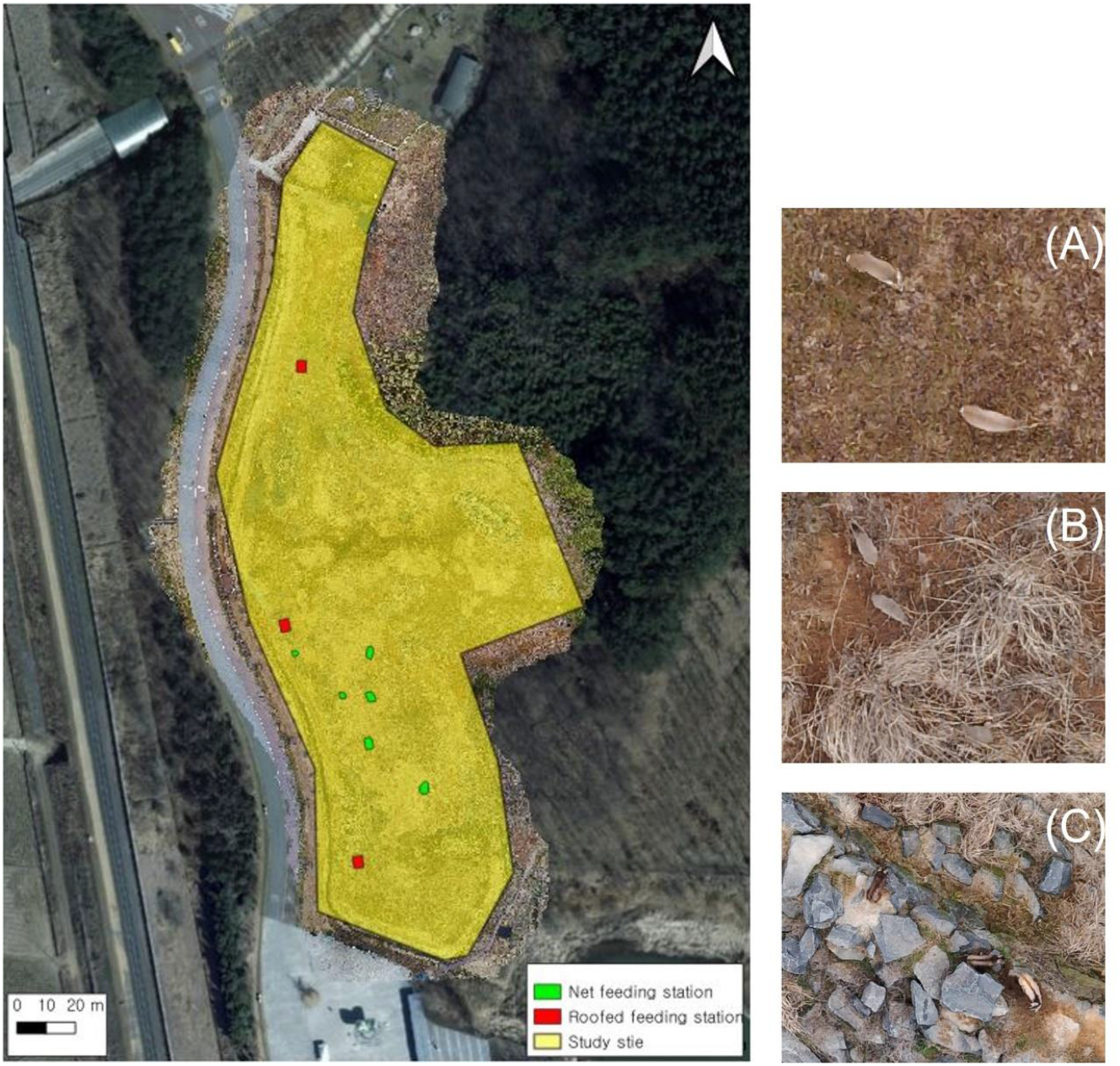
Study area (Deer park) in Seocheon, South Korea. Yellow polygon indicates the study site housing three Korean ungulate species ((A) Roe deer, (B) Water deer, (C) Long-tailed goral). Green and red polygons indicate the different types of feeding station. Basemap sources: Kakao Satellite map(https://map.kakao.com/?URLX=487448&URLY=1504827&URLLEVEL=14&map_type=TYPE_SKYVIEW&map_hybrid=true&SHOWMARK=false)

### 2.2. Data collection

For this study, we utilized two drones: the DJI Matrice 350 RTK, equipped with a Zenmuse H20T camera, and the DJI Mavic Pro 3 Thermal. Both drones simultaneously captured thermal images and regular photographs, with the thermal cameras having a resolution of 640×512 pixels. Prior to capturing orthophotos, we assessed the animals’ reactions by gradually lowering the flight altitude to 10 meters and observed no notable reactions. The minimum flight altitude was set at 30 meters, based on the height of the surrounding canopy. To minimize ground surface radiation, drone flights were conducted near sunrise and sunset. We conducted 17 flights (9 in March and 8 in June), but 2 flights were excluded due to quality issues. We selected March and June for the flights to assess potential seasonal variations in animal visibility and detection. Using DJI Terra software(“DJI Terra,” 2024) on a laptop equipped with an Intel Core™ i7-9857H @ 2.60GHz (12 cores) CPU and a Quadro RTX 3000 GPU, we generated 15 TIR and 15 RGB orthomosaic maps.

### 2.3. Ungulate counts

The ungulate counts from orthomosaic maps were performed using the QGIS pinpoint marker tool. Initially, the counter marked thermal signatures resembling ungulates on the TIR orthomosaic maps (T). These markings were then cross-verified with RGB orthomosaic maps to estimate the observed number of ungulates (TW). The counting error, defined as the difference between the population estimates from the TIR maps and the RGB orthomosaic maps (T-TW), was analyzed in relation to the GSD values to identify the most effective GSD values for accurate population estimation. GSD refers to the real-world size of one pixel in the orthomosaic map and is influenced by flight altitude, camera settings, and sensor resolution. Smaller GSD values (higher resolution) enable more precise detection of features such as thermal signatures of animals, while larger GSD values (lower resolution) can lead to less detailed images and higher counting errors(Fudala and Bialik, 2022; Stone and Davis, 2024). As such, GSD was chosen as a critical variable in this study to assess its impact on the accuracy of animal counts. Detailed results are provided in **Table A1**. A Student’s t-test was applied to compare the average counting error between March and June. We assumed that the counting error would increase as the quality of the maps, indicated by higher GSD values, decreased.

### 2.4. Thermal mark measures for species identification

To evaluate the potential for species identification based on thermal body measurements, we measured the body sizes of thermal signatures from three ungulate species. Specifically, we measured the lengths of four distinct body parts (**Figure 2**) from 149 thermal signatures of Long-tailed goral, roe deer, water deer, and fawn (the young of roe deer or water deer, which are difficult to distinguish between the two species). Species were clearly identified through cross-verification with RGB orthomosaic maps. Thermal signatures that were indistinct, overlapped due to movement, or captured in postures that impeded accurate length measurement were excluded from the analysis. We also classified the postures as standing and lying to account for potential distortions in body measurements.

**Figure 2.**
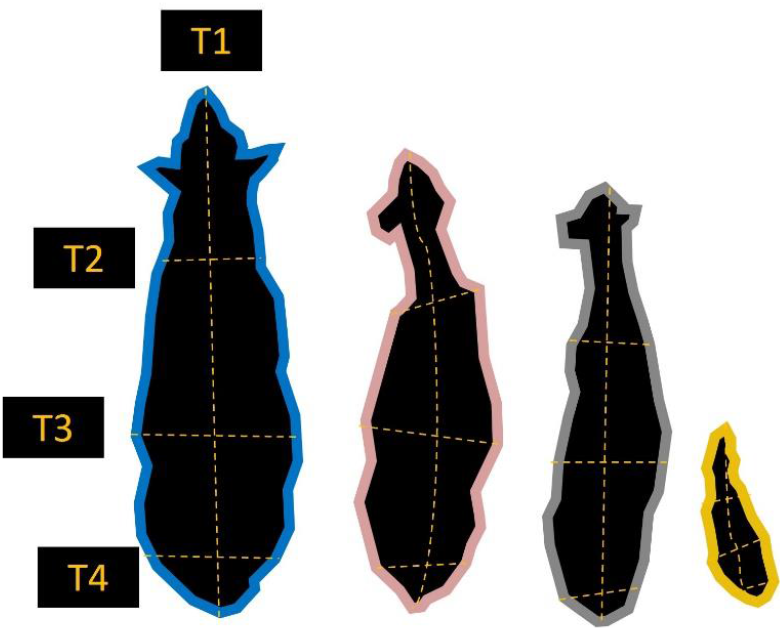
Illustration of thermal mark measurement. T1(Total length from head to tail), T2(Shoulder width), T3(Waist width), T4(Hip width) indicate the length or widths of different body parts of species.

To investigate species-specific differences, we first applied Principal Component Analysis (PCA) to visualize variations in body measurements among the species (Larsen et al., 2023; Ringnér, 2008). We then conducted Multivariate Analysis of Variance (MANOVA) to assess the impact of species and posture on the thermal body measurements. This analysis evaluated differences in the combination of body part lengths and widths across species while considering their interaction (Adams et al., 2013). All statistical analyses and visualizations were performed using R version 4.1.2 (2023)

## 3. Results

### 3.1. Ungulate counts via TIR and RGB orthomosaic maps

The results of the population estimation (**Figure 3**) revealed that the average estimated number of individuals was 25.3 ± 4.2 (range: 18–29) in March and 28.2 ± 9.1 (range: 14–40) in June. The difference in population size between March and June was 4.1 individuals (95% CI: 1.3–6.9), with the T-test showing no clear difference in the average T-TW between the two seasons (p-value = 0.75). We performed regression analysis to quantify the relationship between the total ground sample distance (GSD) and the counting error (**Figure 4**). The estimation error sharply increased when WGSD exceeded 2.5 cm/pix and when TGSD exceeded 7.5 cm/pix.

**Figure 3.**
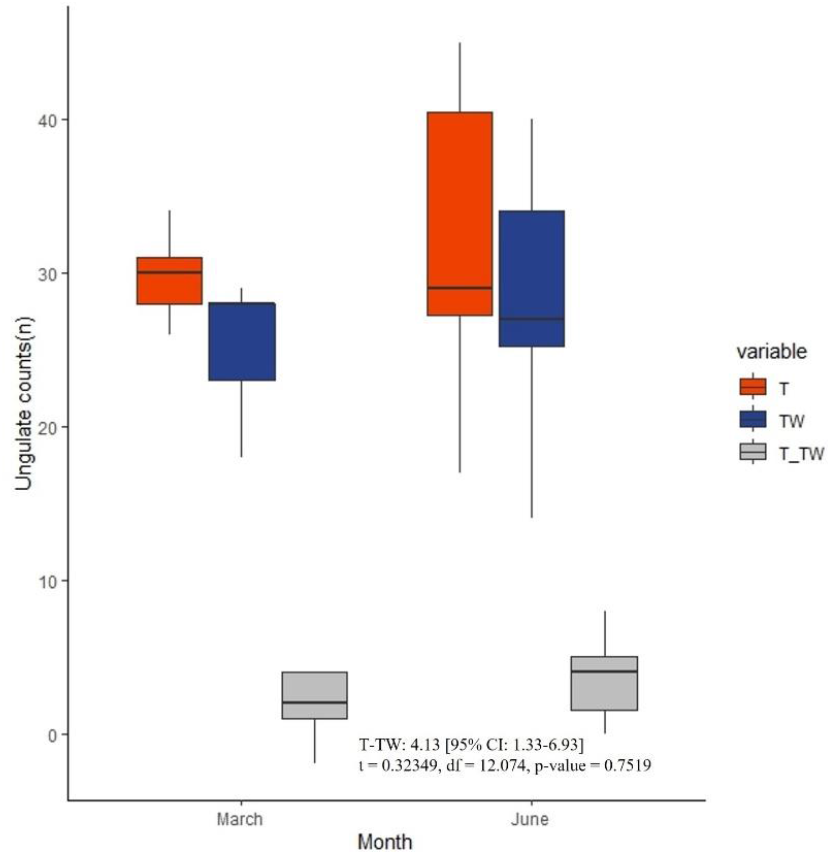
Boxplots illustrating ungulate counts and counting errors for March and June. The plots compare counts derived from TIR orthomosaic maps (T) and Thermal and RGB orthomosaic maps (TW), as well as the counting errors (T-TW).

**Figure 4.**
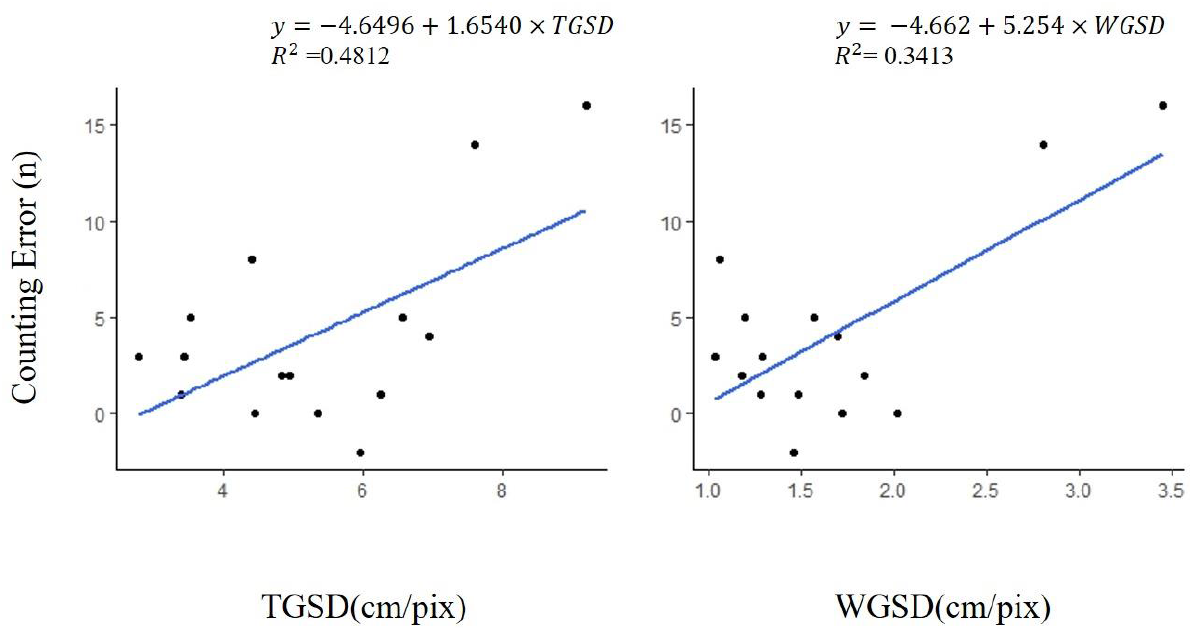
Scatter plots showing the relationship between GSD (cm/pix) and counting error (n), with trend lines fitted using linear regression models. The equations represent the slopes and intercepts of the regression lines. The R^2^ value indicates the proportion of variance in counting error explained by the GSD.

### 3.2. Results of Body Metrics by Species and Posture

We measured the lengths and widths of 149 thermal marks of three ungulate species captured in TIR orthomosaic maps to determine whether these measurements are suitable for species identification. We created boxplots (**Figure 5**) to compare the lengths and widths of thermal marks across species, separating the measurements by posture (lying and standing). The results showed that standing postures provided clearer and more distinguishable measurements compared to lying postures. Among the species, the Long-tailed goral stood out as the largest, with a mean body length of 0.98m and a shoulder width of 0.22m, both surpassing the corresponding values for the other species.”

**Figure 5.**
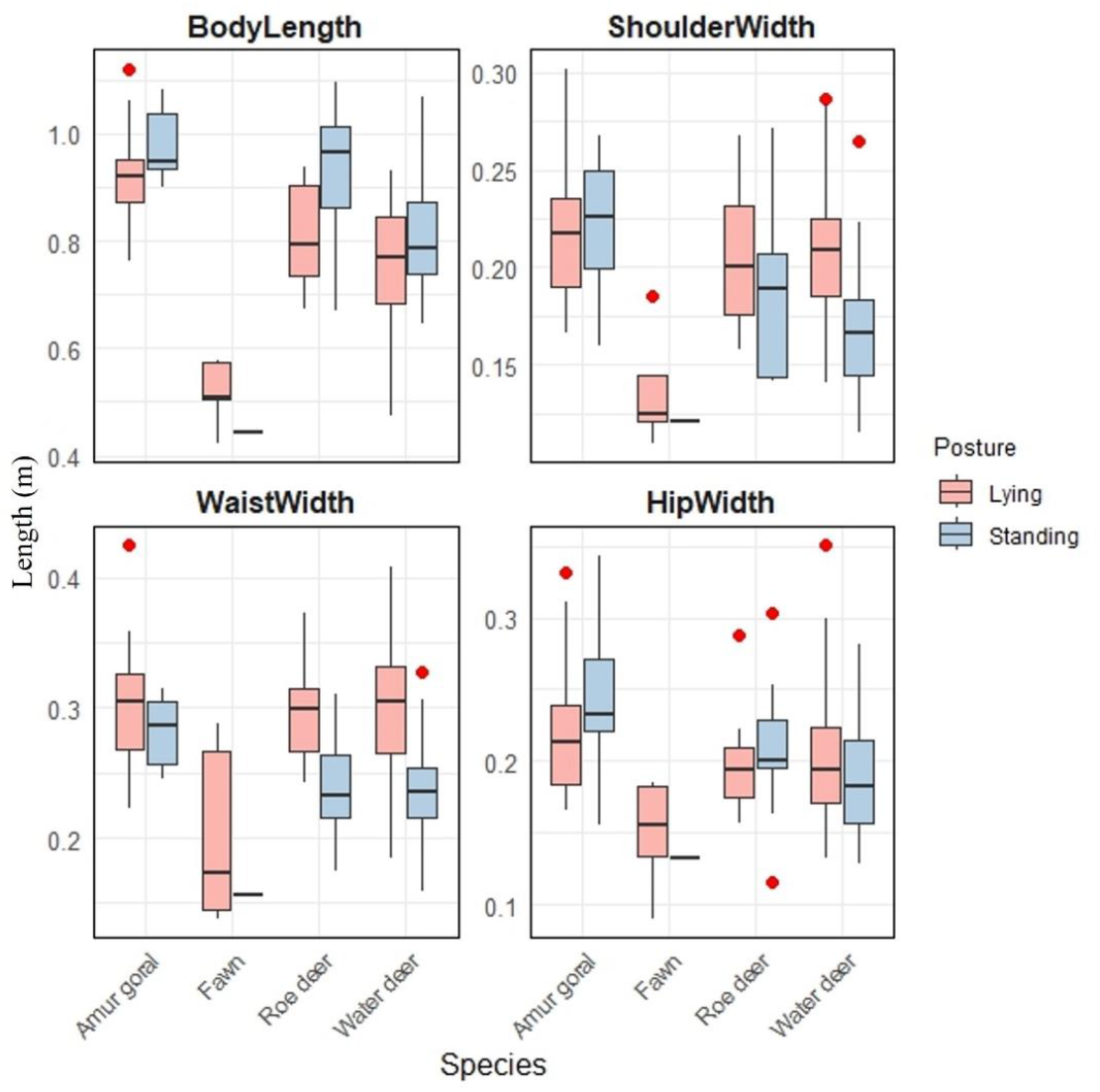
Box plots of thermal mark measures from TIR orthomosaic maps. Boxplots depict the average length and width of four different body part of three Korean ungulate species (Long-tailed (Amur) goral: n = 25; Roe deer: n = 22; Water deer: n = 96; Fawn: n = 6;). Fawn refers to the young of roe deer or water deer, which are difficult to distinguish between the two species. Posture was categorized as lying and standing.

To visualize the differences in body measurement, we conducted principal component analysis (PCA). In the principal component analysis (**Figure 6 (A)**), the first component (PC1) explained 60.13% of the variance, while the second component (PC2) explained an additional 19.5%, cumulatively accounting for 79.6% of the total variance. Based on these results, the first two principal components were selected for further analysis and visualization in the biplot. This component shows strong negative loadings for all four variables: total length (T1: −0.42), shoulder width (T2: −0.52), waist width (T3: −0.52), and hip width (T4: −0.51). PC2 shows a large positive loading for total length (T1: 0.76) and smaller negative loadings for shoulder width (T2: −0.38) and waist width (T3: −0.45), with a modest positive loading for hip width (T4: 0.23). PC2 appears to contrast body length against body width, particularly focusing on the torso area (shoulder and waist widths). This suggests that PC2 captures shape variations, with animals that have longer bodies (high T1) but relatively narrower torso dimensions (T2 and T3) loading strongly on PC2. To sum up, PC1 is primarily a size component, as it indicates proportional increases or decreases across all body measurements. PC2 captures shape differences, with an emphasis on contrasting overall length with torso width, particularly at the shoulder and waist.

**Figure 6.**
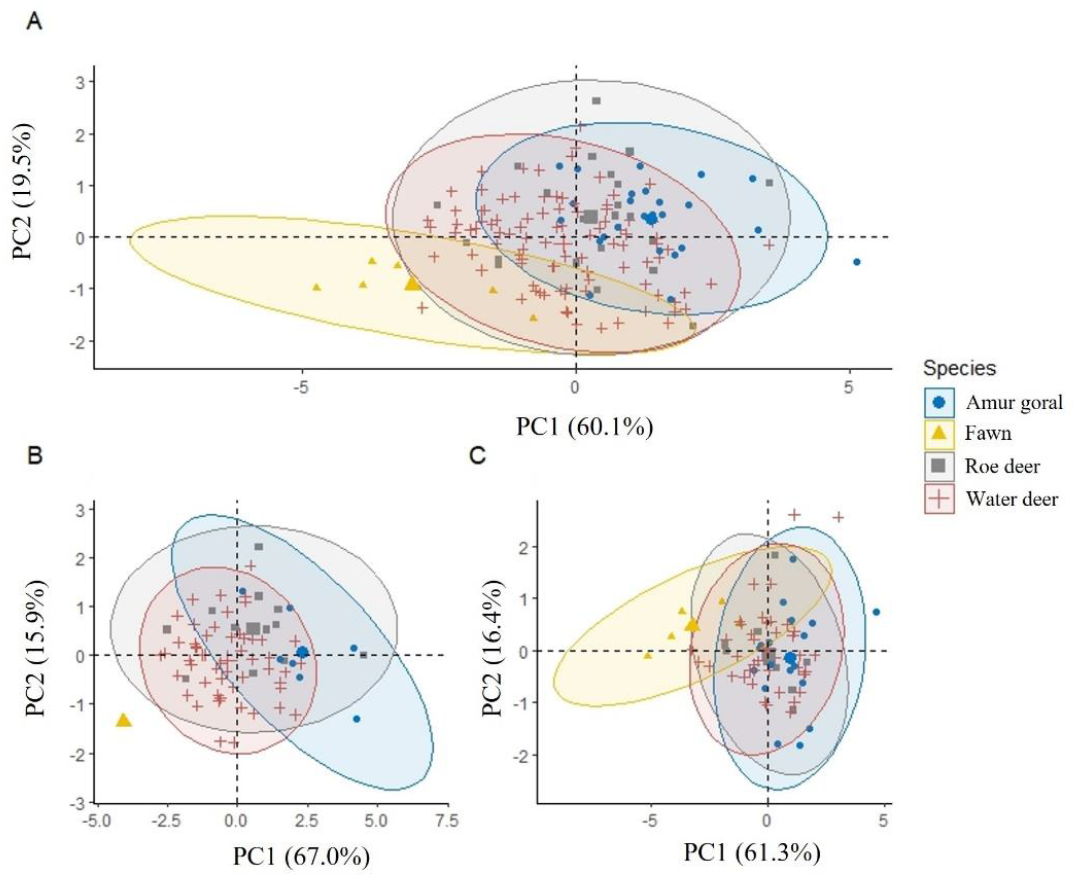
Principal Component Analysis (PCA) ordination of thermal body measurements from TIR orthomosaic maps. (A) PCA combining both standing (n=73) and lying (n=76) postures, (B) PCA for standing posture only, (C) PCA for lying posture only. The first principal component (PC1) accounts for the highest variation among body measurements, while the second principal component (PC2) captures the second highest variation. The body measurements considered include body length, shoulder width, waist width, and hip width of the following species: Long-tailed (Amur) goral (n=25), Roe deer (n=22), Water deer (n=96), and Fawn (n=6). Fawn refers to the young of roe deer or water deer, which are difficult to distinguish from each other.

To assess the effects of species and posture on the species-specific differences in measurements, we used two-way MANOVA (**Table 1**). MANOVA result revealed notable effects of species (P<0.001), indicating substantial differences in body metrics across the various species. However, we also found that the effect of posture was also highly significant (P<0.001) that posture plays a crucial role in the measurement variations. To further assess species-specific differences in body measurements, Canonical Discriminant Analysis (CDA) was conducted. A summary of the canonical coefficients and classification accuracy is provided in **Table 2**. Group means showed that Long-tailed goral had the highest body length and waist width among the species, while Roe deer and Water deer exhibited similar result, except for a longer body length in Roe deer.

**Table 1.**
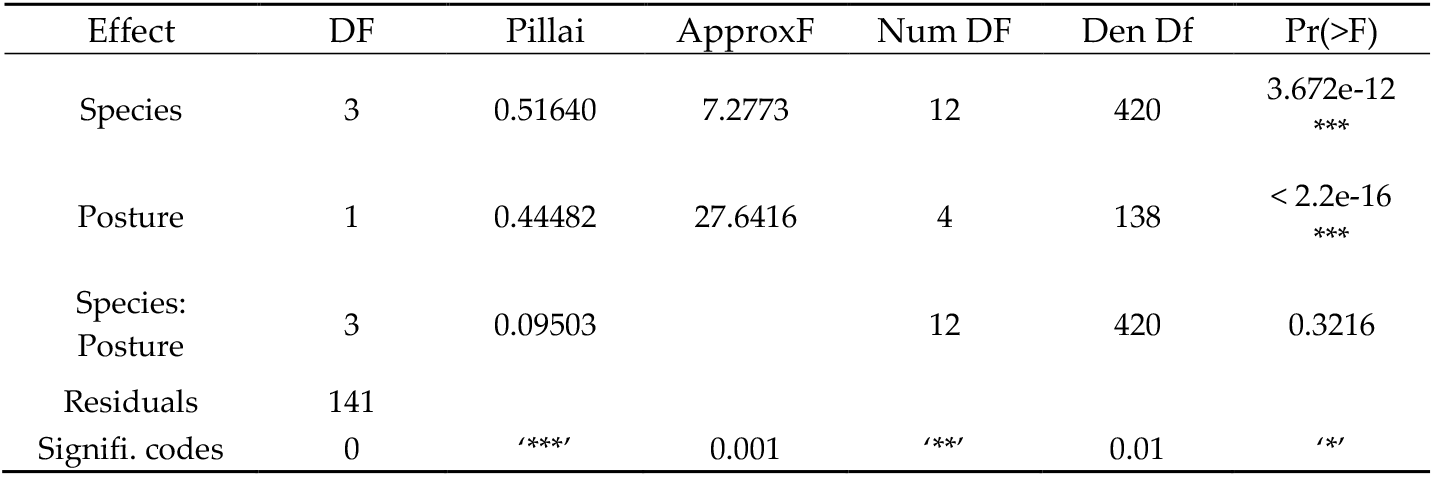
MANOVA Results for Effects of Species and Posture on Body Metrics.

**Table 2.**
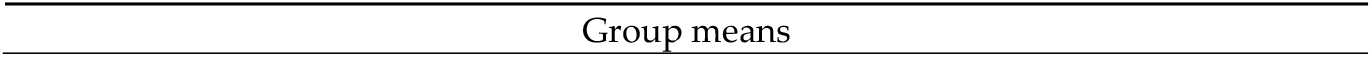
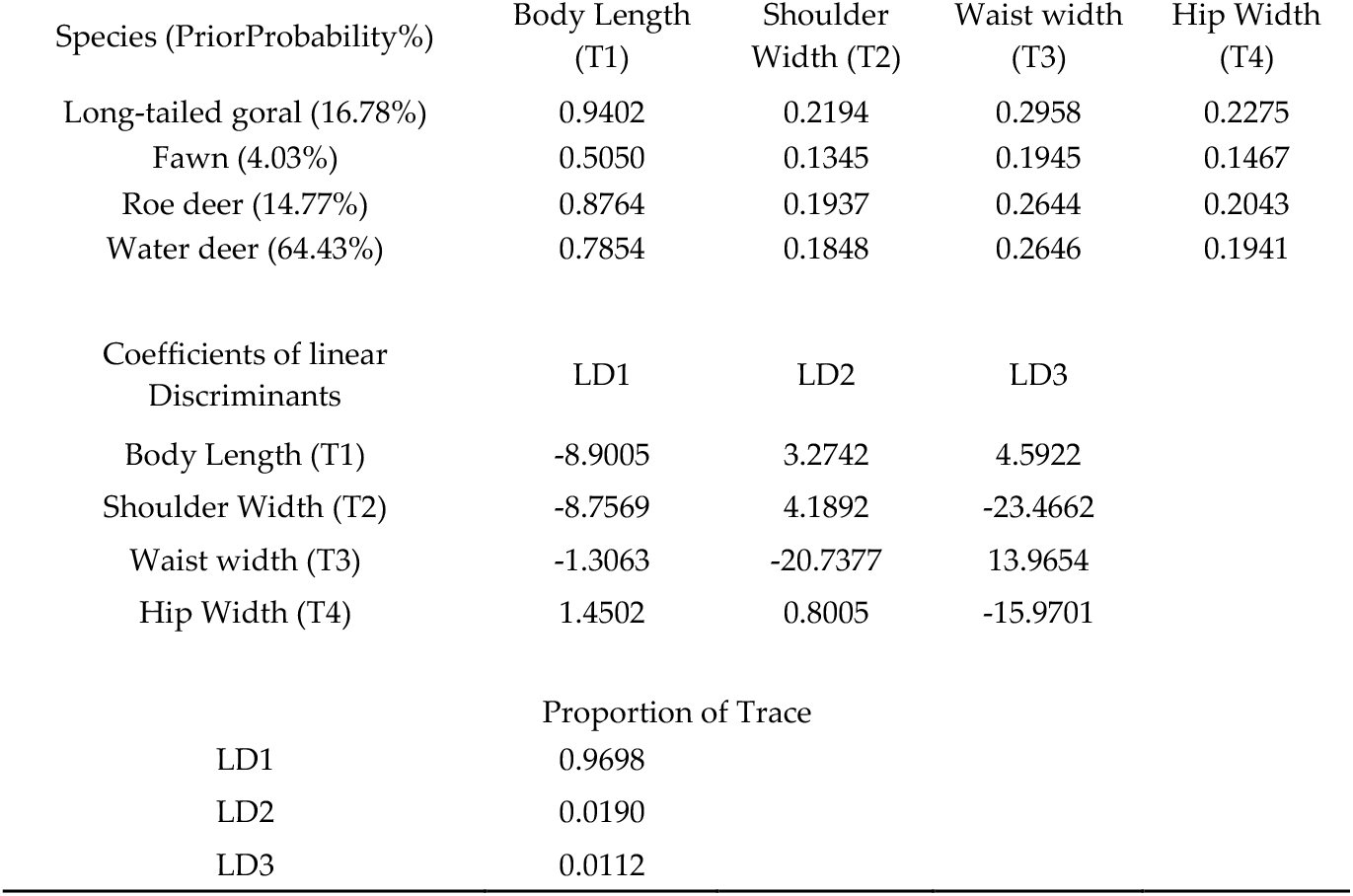
Results of Canonical Discriminant Analysis (CDA) showing group means, coefficients of linear discriminants, and proportion of variance explained by each discriminant function.

| Species (PriorProbability%) | Body Length<br>(T1) | Shoulder<br>Width (T2) | Waist width<br>(T3) | Hip Width<br>(T4) |
| --- | --- | --- | --- | --- |
| Long-tailed goral (16.78%) | 0.9402 | 0.2194 | 0.2958 | 0.2275 |
| Fawn (4.03%) | 0.5050 | 0.1345 | 0.1945 | 0.1467 |
| Roe deer (14.77%) | 0.8764 | 0.1937 | 0.2644 | 0.2043 |
| Water deer (64.43%) | 0.7854 | 0.1848 | 0.2646 | 0.1941 |
| Coefficients of linear<br>Discriminants | LD1 | LD2 | LD3 |  |
| Body Length (T1) | -8.9005 | 3.2742 | 4.5922 |  |
| Shoulder Width (T2) | -8.7569 | 4.1892 | -23.4662 |  |
| Waist width (T3) | -1.3063 | -20.7377 | 13.9654 |  |
| Hip Width (T4) | 1.4502 | 0.8005 | -15.9701 |  |
|  | Proportion of Trace |  |  |  |
| LD1 | 0.9698 |  |  |  |
| LD2 | 0.0190 |  |  |  |
| LD3 | 0.0112 |  |  |  |

To enhance our understanding of the reliability of thermal measurements, we compared the body measurements derived from thermal images (TImeasure) and thermal orthomosaic maps (TOmeasure), which only included standing posture measurements. The detailed results of this comparision are shown in **Table A2 (Appendix B)**. We collected 12 thermal images of three ungulate species taken from a perpendicular angle at approximately 70m above the ground. The comparison revealed no significant differences between the two methods (p-values > 0.05), indicating that TOmeasure effectively captures the actual physical dimensions of the ungulates, supporting its robustness as a method for measuring body metrics. Specifically, the t-test results for body length (p=0.991), shoulder width (p=0.967), waist width (p=0.6301), and hip width (p=0.9119) all showed no strong differences.

## 4. Discussion

In this study, we evaluated the feasibility of using TIR orthomosaic maps for counting three Korean ungulate species. We successfully orthorectified 15 maps, each of RGB and TIR imagery. The counting errors observed in March were primarily due to visibility bias resulting from individuals hiding under feeding stations, which made them difficult to detect using orthomosaic maps. Additionally, other factors that may have contributed to inaccuracies in population estimation, including animals that were moving faster than the drone flight speed and those that could not be cross-verified with RGB orthomosaic maps due to obstructions like grown plants, bushes, or reeds. The increase in population size in June is likely attributed to the appearance of offspring of water deer and roe deer, which were often challenging to detect in the maps. Furthermore, seasonal flushing, new shoot, and ground surface radiation reduced visibility, contributing to the larger confidence intervals and broader ranges of T and TW observed in June.

Linear regression analyses were performed to investigate the relationship between GSD and the counting error using two types of GSD measurements: TGSD and WGSD. TGSD, which typically has a higher GSD due to the lower quality of thermal images, showed a positive relationship with counting error. However, the model explained only 34.1% of the variance in counting error, suggesting that while TGSD affects counting error, other factors may also play a role. In contrast, WGSD, which has a lower GSD due to the higher quality of RGB orthomosaic maps, demonstrated a stronger relationship with counting error. The coefficient for WGSD (3.5cm/pix) was higher than that for TGSD, and the model explains 48.1% of the variance in counting error, with an R^2^ value of 0.48. A significant increase in counting errors was observed when TGSD exceeded 7.5cm/pix and WGSD exceeded 2.5cm/pix. These findings suggest potential threshold values for GSD that could be useful as reference points in future study designs. Specifically, maintaining TGSD below 7cm/pix and WGSD below 2.5cm/pix may help reduce counting errors and improve the accuracy of orthomosaic map-based analyses.

While we successfully counted the animals in the enclosure, it is important to acknowledge that this setup may not accurately reflect the challenges encountered in wild populations. Wild animals often exhibit different behaviors and movement patterns compared to those in controlled environments, which can influence detection rates and the accuracy of population estimates(Samuel et al., 1992; Willson et al., 2011). Factors such as habitat complexity(Melbourne, 1999; Zhang et al., 2021), availability of resources(Converse et al., 2006), and human disturbances(Varman and Sukumar, 1995) can significantly affect wildlife monitoring. In particular, wild ungulates species in high-forested areas are often underestimated in previous studies(Cukor et al., 2022; Peña-Carmona et al., 2024; Tóth and Katona, 2024). Therefore, the findings from this study may not be directly applicable to wild populations without further validation. In addition to the factors previously discussed, it is important to note that Korean ungulates are generally solitary outside of the breeding season, which poses challenges for population estimation throughout the year. Given the increased visibility of these animals during winter due to leafless trees and cooler temperatures, as well as the breeding season typically occurring in autumn and winter, we recommend that future studies focus on these seasons. Conducting research during autumn and winter will likely enhance detection rates and provide more accurate population estimates, ultimately improving our understanding of these species in their natural habitats.

We found thermal mark measurements from TIR orthomosaic maps ineffective for species identification, as variations in posture significantly impacted body metrics, increasing the likelihood of misclassification. Specifically, when measuring body length in a lying position, there was significant overlap between species, as the width of the ungulates appeared broader due to compression of the back and pelvis when lying down. The MANOVA results also supported that posture had a strong effect on body metrics, highlighting the challenge of using thermal mark measurements for species identification. This challenge is not unique to our study; similar difficulties have been observed in other research, such as the measurement of body size in Australian fur seals(Allan et al., 2019), where postural sag in the lateral thorax when lying on rocks led to overestimation of body measurements. Moreover, in studies aimed at estimating body volumes, like those involving pinnipeds(Stone and Davis, 2024), movement between images and body posture have been shown to hinder accurate measurements, leading to less successful estimates. Although species differentiation was somewhat clearer when measuring in a standing position in the present study, relying solely on length measurements for species identification remains a practical challenge.

However, we did not find the use of thermal orthomosaic maps (TO) for measuring body size to be an inappropriate method. The comparison between thermal images (TI) and TOmeasure showed that TOmeasure is capable of accurately capturing the physical dimensions of the ungulates. This suggests that, while species identification remains a challenge, TOmeasure can still be a reliable tool for estimating body size and monitoring populations. The lack of existing data on average body metrics for these ungulate spcies—particularly measurements taken at perpendicular angels—may have contributed to the difficulty in species differentiation. Therefore, further data collection, including additional thermal image and orthomosaic map analyses across different environments, would likely reinforce our findings and improve the accuracy of this method for species identification.

## Ethics approval

This study was entirely observational and did not involve any direct interactions with or experiments on animals, so ethics approval was not required.

## Author Contributions

Conceptualization, J.K. and D.W.; methodology, J.K. and D.W.; software, J.K.; validation, J.K. and D.W.; formal analysis, J.K. and D.W.; investigation, J.K. and D.W.; resources, J.K. and D.W.; data curation, J.K.; writing—original draft preparation, J.K.; writing—review and editing, J.K. and D.W.; visualization, J.K.; supervision, D.W.; project administration, D.W..; funding acquisition, D.W. All authors have read and agreed to the published version of the manuscript.

## Data Availability Statement

Data may be shared by contacting the corresponding author.

## Acknowledgments

We would like to thank the Department of Animal & Exhibition in National Institute of Ecology for granting permission to take drone pictures at deer park, which greatly contributed to the success of this study.

## Appendix A. Ungulate counts

TIR count is the number of counted ungulates from TIR orthomosaic maps. TW is the actual number of ungulate individuals which were cross-verified with RGB orthomosaic maps. T-TW is the differences between the estimated population numbers in the thermal orthomosaic maps and the RGB orthomosaic maps. NIE test2-5, 2-6 are removed due to the low visibility from RGB orthomosaic maps.

**Table A1.**
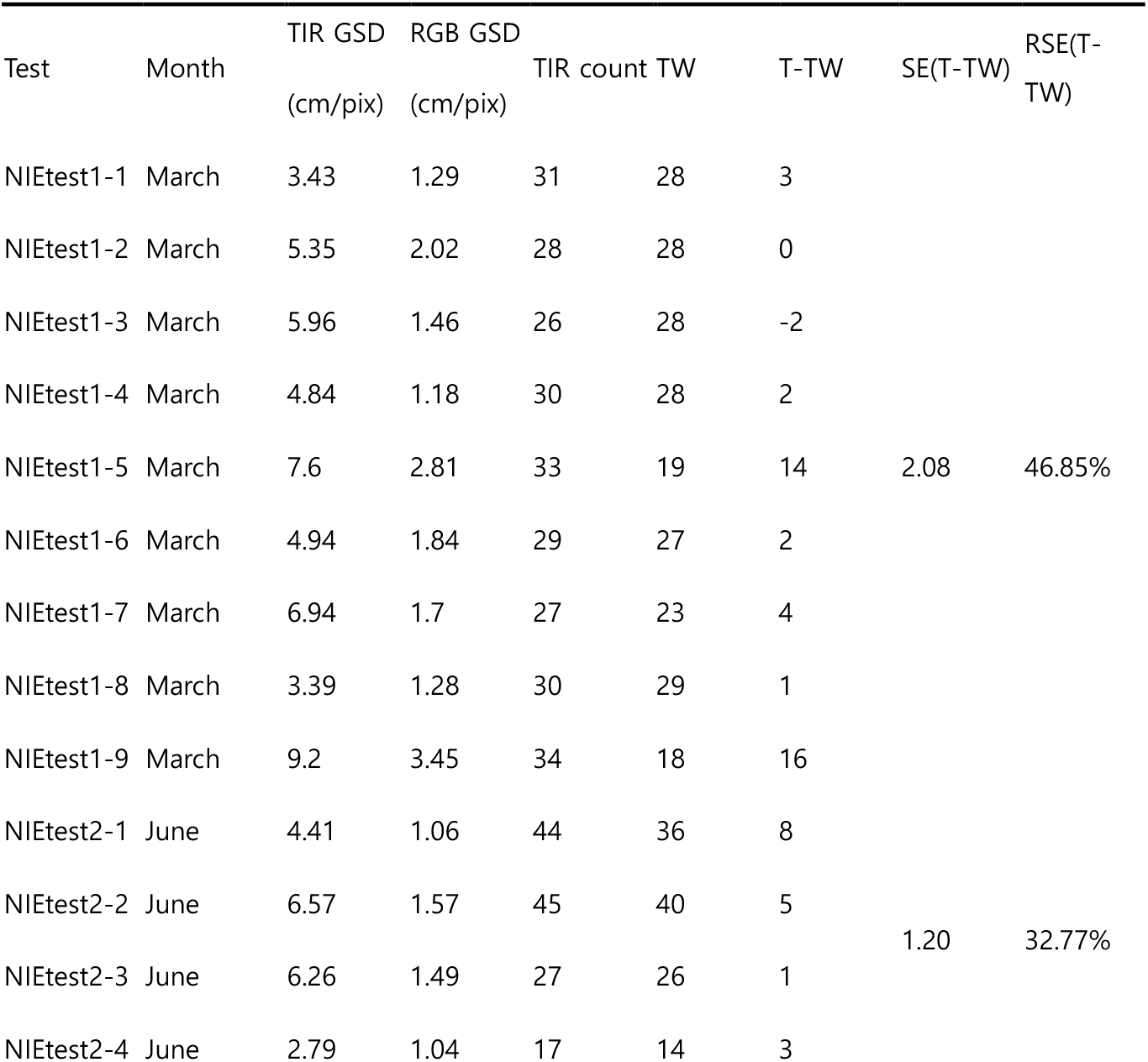
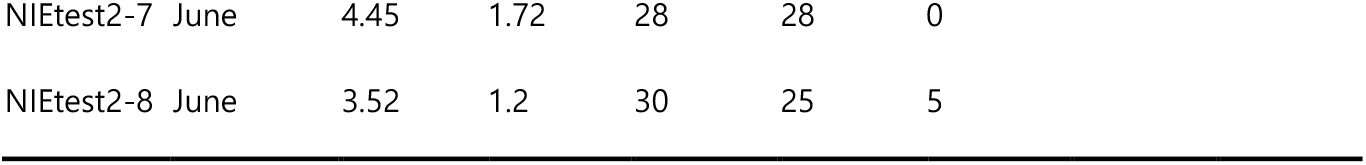
Drone flight specifications and ungulate count results.

## Appendix B. Comparison of body metrics for ungulate species derived from Thermal Orthomosaic maps (TOmeasure) and Thermal Images (TImeasure)

This table presents the average body length, shoulder width, waist width, and hip width measurements for four ungulate species: Long-tailed goral, Roe deer, Water deer, and Fawn(roe deer or water deer). The measurements obtained from thermal orthomosaic maps (TOmeasure) are compared to those obtained from thermal images (TImeasure).

P-values indicate the statistical significance of differences between the two measurement methods. All p-values are significantly greater than 0.05, indicating no statistically significant difference between the mean values of the two measurements for any of the body metrics

**Table A2.** Comparison of body metrics for ungulate species derived from Thermal Orthomosaic maps (TOmeasure) and Thermal Images (TImeasure)

|  | Thermal image measures (TI) |  |  |  | Thermal orthomosaic measures (TO) |  |  |  |
| --- | --- | --- | --- | --- | --- | --- | --- | --- |
| Body parts | Body length | Shoulder width | Waist width | Hip width | Body length | Shoulder width | Waist width | Hip width |
| Species |  |  |  |  |  |  |  |  |
| Long-tailed goral | 1.015 | 0.246 | 0.286 | 0.225 | 0.982 | 0.222 | 0.281 | 0.245 |
| Roe deer | 0.89 | 0.157 | 0.25 | 0.2 | 0.933 | 0.185 | 0.237 | 0.209 |
| Water deer | 0.797 | 0.157 | 0.239 | 0.187 | 0.804 | 0.167 | 0.236 | 0.190 |
| Fawn | 0.452 | 0.129 | 0.198 | 0.15 | 0.442 | 0.121 | 0.156 | 0.132 |
| P-value | 0.9916 | 0.9670 | 0.6301 | 0.9119 |  |  |  |  |

P-values indicate the statistical significance of differences between the two measurement methods. All p-values are significantly greater than 0.05, indicating no statistically significant difference between the mean values of the two measurements for any of the body metrics

## References

Adams, D.C., Rohlf, F.J., Slice, D.E., 2013. A field comes of age: geometric morphometrics in the 21st century. Hystrix, the Italian Journal of Mammalogy 24. 10.4404/hystrix-24.1-6283

Allan, B.M., Ierodiaconou, D., Hoskins, A.J., Arnould, J.P.Y., 2019. A Rapid UAV Method for Assessing Body Condition in Fur Seals. Drones 3, 24. 10.3390/drones3010024

Amos, M., Baxter, G., Finch, N., Lisle, A., Murray, P., 2014. “I just want to count them! Considerations when choosing a deer population monitoring method.” Wildlife Biology 20, 362–370. 10.2981/wlb.00080

Beaver, J.T., Baldwin, R.W., Messinger, M., Newbolt, C.H., Ditchkoff, S.S., Silman, M.R., 2020. Evaluating the Use of Drones Equipped with Thermal Sensors as an Effective Method for Estimating Wildlife. Wildl. Soc. Bull. 44, 434–443. 10.1002/wsb.1090

Converse, S.J., Block, W.M., White, G.C., 2006. Small mammal population and habitat responses to forest thinning and prescribed fire. Forest Ecology and Management 228, 263–273. 10.1016/j.foreco.2006.03.006

Cukor, J., Havránek, F., Sokolov, S., Skoták, V., Hambálková, L., Ševčík, R., Vacek, Z., Nurseitov, D., 2022. Estimation of ungulate population density in Kazakhstan: Case study from foothill ecosystems. J. For. Sci. 68, 452–458. 10.17221/98/2022-JFS

Davis, S., n.d. Evaluating the Use of Drones to Estimate Deer Density and Count Wildlife Trails in Bath Nature Preserve. OhioLINK Electronic Theses and Dissertations Center. DJI Terra, 2024.

Dundas, S.J., Vardanega, M., O’Brien, P., McLeod, S.R., 2021. Quantifying Waterfowl Numbers: Comparison of Drone and Ground-Based Survey Methods for Surveying Waterfowl on Artificial Waterbodies. Drones 5, 5. 10.3390/drones5010005

Ezat, M.A., Fritsch, C.J., Downs, C.T., 2018. Use of an unmanned aerial vehicle (drone) to survey Nile crocodile populations: A case study at Lake Nyamithi, Ndumo game reserve, South Africa. Biological Conservation 223, 76–81. 10.1016/j.biocon.2018.04.032

Fudala, K., Bialik, R.J., 2022. The use of drone-based aerial photogrammetry in population monitoring of Southern Giant Petrels in ASMA 1, King George Island, maritime Antarctica. Global Ecology and Conservation 33, e01990. 10.1016/j.gecco.2021.e01990

Greenhorn, J.E., Sadowski, C., Rodgers, J.A., Bowman, J., 2024. The use of orthoimagery and stereoscopic aerial imagery to identify muskrat (Ondatra zibethicus) houses. Wildlife Society Bulletin e1519. 10.1002/wsb.1519

Gu, W., Swihart, R.K., 2004. Absent or undetected? Effects of non-detection of species occurrence on wildlife–habitat models. Biological Conservation 116, 195–203. 10.1016/s0006-3207(03)00190-3

Inman, V.L., Kingsford, R.T., Chase, M.J., Leggett, K.E.A., 2019. Drone-based effective counting and ageing of hippopotamus (Hippopotamus amphibius) in the Okavango Delta in Botswana. PLoS ONE 14, e0219652. 10.1371/journal.pone.0219652

Jang, J.E., Kim, N.H., Lim, S., Kim, K.Y., Lee, H.J., Park, Y.C., 2020. Genetic integrity and individual identification-based population size estimate of the endangered long-tailed goral, Naemorhedus caudatus from Seoraksan National Park in South Korea, based on a non-invasive genetic approach. Animal Cells and Systems 24, 171–179. 10.1080/19768354.2020.1784273

Kim, B.-J., Oh, D.-H., Chun, S.-H., Lee, S.-D., 2011. Distribution, density, and habitat use of the Korean water deer (Hydropotes inermis argyropus) in Korea. Landscape Ecol Eng 7, 291–297. 10.1007/s11355-010-0127-y

Larsen, H.L., Møller-Lassesen, K., Enevoldsen, E.M.E., Madsen, S.B., Obsen, M.T., Povlsen, P., Bruhn, D., Pertoldi, C., Pagh, S., 2023. Drone with Mounted Thermal Infrared Cameras for Monitoring Terrestrial Mammals. Drones 7, 680. 10.3390/drones7110680

Mayer, M., Furuhovde, E., Nordli, K., Myriam Ausilio, G., Wabakken, P., Eriksen, A., Evans, A.L., Mathisen, K.M., Zimmermann, B., 2024. Monitoring GPS‐collared moose by ground versus drone approaches: efficiency and disturbance effects. Wildlife Biology e01213. 10.1002/wlb3.01213

McCarthy, E.D., Martin, J.M., Boer, M.M., Welbergen, J.A., 2022. Ground-based counting methods underestimate true numbers of a threatened colonial mammal: an evaluation using drone-based thermal surveys as a reference. Wildlife Res. 50, 484–493. 10.1071/WR21120

Melbourne, B.A., 1999. Bias in the effect of habitat structure on pitfall traps: An experimental evaluation. Australian Journal of Ecology 24, 228–239. 10.1046/j.1442-9993.1999.00967.x

Mirka, B., Stow, D.A., Paulus, G., Loerch, A.C., Coulter, L.L., An, L., Lewison, R.L., Pflüger, L.S., 2022. Evaluation of thermal infrared imaging from uninhabited aerial vehicles for arboreal wildlife surveillance. Environ Monit Assess 194, 512. 10.1007/s10661-022-10152-2

Peña-Carmona, G., Escobar-González, M., Dobbins, M.T., Conejero, C., Valldeperes, M., Lavín, S., Pérez, J.M., López-Olvera, J.R., López-Martín, J.M., Serrano, E., 2024. Direct counts underestimate mountain ungulate population size. 10.21203/rs.3.rs-4009600/v1

Povlsen, P., Bruhn, D., Pertoldi, C., Pagh, S., 2023. A Novel Scouring Method to Monitor Nocturnal Mammals Using Uncrewed Aerial Vehicles and Thermal Cameras—A Comparison to Line Transect Spotlight Counts. Drones 7, 661. 10.3390/drones7110661

R Core Team, 2023. R: A Language and Environment for Statistical Computing.

Ringnér, M., 2008. What is principal component analysis? Nat Biotechnol 26, 303–304. 10.1038/nbt0308-303

Samuel, M.D., Steinhorst, R.K., Garton, E.O., Unsworth, J.W., 1992. Estimation of Wildlife Population Ratios Incorporating Survey Design and Visibility Bias. The Journal of Wildlife Management 56, 718. 10.2307/3809465

Stone, T.C., Davis, K.J., 2024. Using unmanned aerial vehicles to estimate body volume at scale for ecological monitoring. Methods Ecol Evol 2041–210X.14457. 10.1111/2041-210X.14457

Tóth, G., Katona, K., 2024. Comparison of Population Density Estimation Methods for Roe Deer (Capreolus capreolus). Diversity 16, 500. 10.3390/d16080500

Varela-Jaramillo, A., Rivas-Torres, G., Guayasamin, J.M., Steinfartz, S., MacLeod, A., 2023. A pilot study to estimate the population size of endangered Galápagos marine iguanas using drones. Front Zool 20, 4. 10.1186/s12983-022-00478-5

Varman, K.S., Sukumar, R., 1995. The line transect method for estimating densities of large mammals in a tropical deciduous forest: An evaluation of models and field experiments. J. Biosci. 20, 273–287. 10.1007/BF02703274

Willson, J.D., Winne, C.T., Todd, B.D., 2011. Ecological and methodological factors affecting detectability and population estimation in elusive species. J Wildl Manag 75, 36–45. 10.1002/jwmg.15

Witczuk, J., Pagacz, S., Zmarz, A., Cypel, M., 2018. Exploring the feasibility of unmanned aerial vehicles and thermal imaging for ungulate surveys in forests - preliminary results. International Journal of Remote Sensing 39, 5504–5521. 10.1080/01431161.2017.1390621

Zhang, W., Sheldon, B.C., Grenyer, R., Gaston, K.J., 2021. Habitat change and biased sampling influence estimation of diversity trends. Current Biology 31, 3656-3662.e3. 10.1016/j.cub.2021.05.066

